# Activity-dependent regulation of vascular cholesterol metabolism acts as a negative feedback mechanism for neurovascular coupling

**DOI:** 10.1101/2024.02.23.581685

**Authors:** CP Profaci, KL Foreman, L Spieth, V Coelho-Santos, SA Berghoff, JT Fontaine, DA Jeffrey, SP Palecek, EV Shusta, G Saher, AY Shih, F Dabertrand, R Daneman

**Affiliations:** Department of Pharmacology, University of California, San Diego. La Jolla, CA, USA; Department of Neurosciences, University of California, San Diego. La Jolla, CA, USA; Department of Chemical and Biological Engineering, University of Wisconsin-Madison. Madison, WI, USA; Department of Neurogenetics, Max Planck Institute for Multidisciplinary Sciences, Göttingen, Germany; Center for Developmental Biology and Regenerative Medicine, Seattle Children’s Research Institute. Seattle, WA, USA; Department of Pediatrics, University of Washington. Seattle, WA, USA; Department of Bioengineering, University of Washington. Seattle, WA, USA; Institute of Neuronal Cell Biology, Technical University Munich. Munich, Germany; German Center for Neurodegenerative Diseases (DZNE). Munich, Germany; Department of Anesthesiology, University of Colorado Anschutz Medical Campus. Aurora, CO, USA; Department of Pharmacology, University of Colorado Anschutz Medical Campus. Aurora, CO, USA

## Abstract

Brain function is dependent on a continuous supply of bloodborne oxygen and nutrients. Because neurons require a greater supply of oxygen and nutrients when active, there is increased local blood flow following neuronal activity. The underlying mechanisms of this hyperemia are termed neurovascular coupling (NVC). Many complex processes contribute to NVC, and there is still much unknown about how vascular physiology adapts to changes in neuronal activity and blood flow. Here we show that neuronal activity increases brain endothelial expression of genes related to cholesterol synthesis and uptake *in vivo*, and that shear stress is sufficient for upregulation of these genes *in vitro*. We previously found that treatment with PLX5622 induces upregulation of the same cassette of cholesterol-related genes in brain endothelial cells. In the present study, we find that increasing brain endothelial cholesterol synthesis and/or uptake, either with PLX5622 or targeted AAV-mediated expression of LDLR, inhibits brain arteriole dilation in response to capillary K^+^ stimulation, and this deficit is rescued by cholesterol depletion. Together, these data suggest that neuronal activity regulates brain endothelial cholesterol, which in turn blocks endothelial retrograde signaling and vasodilation, thus acting as a negative feedback mechanism for NVC.

## Introduction

A constant supply of bloodborne oxygen and nutrients is required for proper brain function, and even a brief lapse in cerebral blood supply leads to catastrophic consequences for brain health. Because neurons require a greater supply of oxygen and nutrients when active, there is increased local blood flow immediately following neuronal activity, a phenomenon observed as early as 1890^1–3^. More than a century later, there is much known—and still much unknown—about the cellular and molecular mechanisms of neurovascular coupling (NVC). Studies have implicated several vasoactive ions and molecules in NVC, including potassium ions (K^+^), nitric oxide, prostaglandin, neuropeptide Y, and vasoactive intestinal peptide^4–6^. These signals ultimately lead to the relaxation of mural cells, causing dilation of underlying blood vessels and increased blood flow^7–11^. It was long thought that endothelial cells play a passive role in this process, but recent work has uncovered that endothelial cells are indeed active participants in NVC^12–14^. There remains much unknown about endothelial participation in and regulation of NVC, including how endothelial cells adapt to increases in neuronal activity and changes in blood flow.

One of the primary ways in which endothelial cells directly participate in NVC is through K^+^ signaling. Neuronal activity increases the extracellular K^+^ concentration, which activates abluminal K_IR_2.1 channels on endothelial cells and smooth muscle cells (SMCs)^13,15^. Capillary K^+^ activation causes endothelial cell hyperpolarization, and this current travels upstream—likely via endothelial gap junctions— leading to arteriole-SMC signaling, SMC relaxation, and arteriole dilation^13,16^. This process allows capillary endothelial cells to sense highly localized changes in neuronal activity and send retrograde signals to the arterioles to alter local blood flow. Interestingly, a collection of studies has identified lipids as mediators of K_IR_ activity. Endothelial depletion of phosphatidylinositol 4,5-biphosphate (PIP_2_), a minor phospholipid component of cell membranes, has been shown to reduce endothelial K_IR_ current, inhibiting capillary K^+^ stimulation-mediated hyperemia and rendering resistance arteries less sensitive to laminar flow^17–19^. Cholesterol has also been shown to modulate K_IR_ current in various cell lines: an increase in cellular cholesterol dampens K_IR_ current, while depleting cholesterol or replacing it with an optical isomer increases current^20,21^. Furthermore, endothelial K_IR_ dysfunction underlies deficits in flow-induced vasodilation in obese mice and humans^22^. However, very little is known about whether endothelial cholesterol levels are dynamically regulated in endothelial cells in health and what functional role this might play in NVC or other vascular physiologies.

Here we show that increasing neuronal activity upregulates brain endothelial expression of cholesterol synthesis enzymes, cholesterol synthesis regulators, and the cholesterol uptake receptor low-density lipoprotein receptor (LDLR). Conversely, silencing neuronal activity has the opposite effect, decreasing expression of these cholesterol-related genes in brain endothelial cells *in vivo*. We further report that shear stress is sufficient to upregulate the cholesterol gene cassette in brain endothelial-like cells *in vitro*, suggesting a role for blood flow in mediating the activity-dependent regulation of brain endothelial cholesterol metabolism. Previously, we found that PLX5622, a drug used to deplete microglia, induces the upregulation of cholesterol synthesis enzymes and uptake receptor specifically in brain endothelial cells in a microglia depletion-independent manner^23^. We therefore use PLX5622 to pharmacologically target central nervous system (CNS) endothelial synthesis and uptake pathways to probe the physiological consequences of changes in brain endothelial cholesterol. We find that altering expression of this cassette of cholesterol synthesis and uptake genes does not affect overall brain cholesterol levels, suggesting that activity-dependent changes in brain endothelial cholesterol metabolism play a local role within endothelial cells. Indeed, we find here that increased endothelial cholesterol, induced either by PLX5622 administration or brain endothelial overexpression of LDLR, suppresses arteriole dilation in response to K^+^ capillary stimulation. Thus, increasing endothelial cholesterol inhibits capillaries from signaling to upstream vascular segments to induce increased blood flow after local neuronal activity. Together, these data illuminate a novel role for neuronal activity in regulating vascular physiology and suggest that brain endothelial cholesterol can act as a negative feedback mechanism for an important aspect of NVC.

## Results

### Neuronal activity regulates brain endothelial cholesterol synthesis in vivo

We previously used chemogenetics to identify how neuronal activity influences brain endothelial cells, performing endothelial transcriptomics under conditions of normal, activated, and silenced neuronal activity. We found that neuronal activity regulates the expression of blood-brain barrier efflux transporters^24^. Remarkably, in continued analysis of this dataset, we found a clear pattern of bidirectional modulation of cholesterol pathway gene expression by neuronal activity: activation of glutamatergic neurons leads to overall higher brain endothelial expression of several genes involved in cholesterol synthesis (*Hmgcr*, *Pmvk*, *Mvd*, *Fdps*, *Sqle*, *Cyp51*, *Msmo1*, *Nsdhl*, *Hsd17b7*, *Sc5d*), synthesis regulation (*Srebf2*), and uptake (*Ldlr*), while silencing glutamatergic neural activity has the opposite effect (**Fig 1A**). Expression of the cholesterol efflux transporter, *Abca1,* trended in the opposite direction from genes coding for synthesis and uptake machinery (**Fig 1A**). We validated these results by measuring LDLR expression in brain endothelial cells from *CamKII*α*-*tTA; TRE-hM3Dq (DREADDs^activating^) and littermate controls after clozapine-N-oxide (CNO) injection. Increased neuronal activity increased the percentage of CD31+ vascular length that was also LDLR+ (p=0.0033) (**Fig 1B**). These data suggest that neuronal activity upregulates brain endothelial cholesterol synthesis and uptake while inhibiting cholesterol efflux *in vivo*.

**Figure 1.**
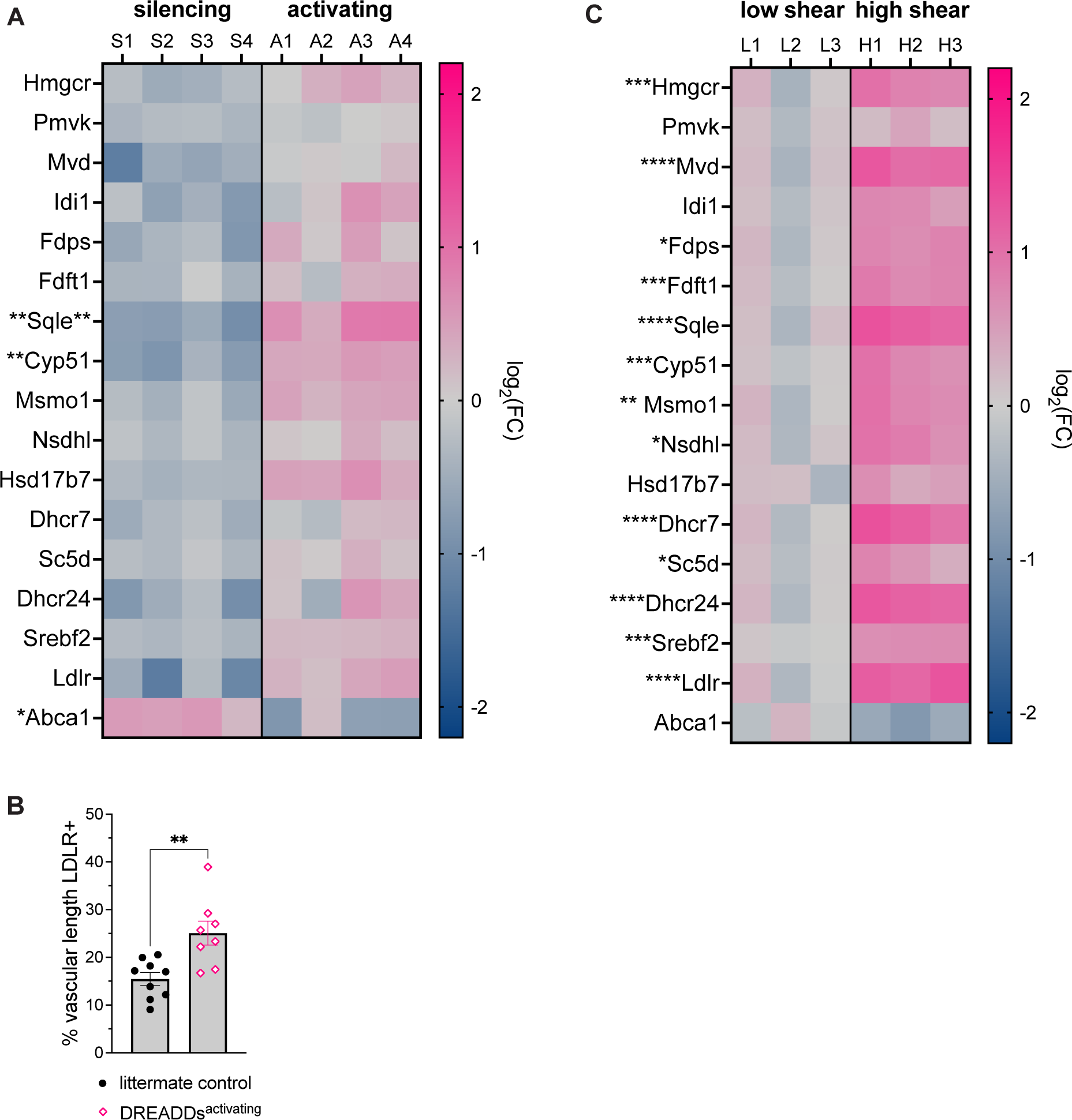
Neuronal activity and shear stress regulate brain endothelial cholesterol synthesis and uptake. (A) Heat map of cholesterol gene expression in brain endothelial cells after neuronal activating (A) or silencing (S) (Pulido et al., 2020). DREADDs^activating^ mice, DREADDs^silencing^ mice, and respective littermate controls were injected with CNO. Bulk RNA sequencing was performed on isolated endothelial cells. Heat map color scale indicates log_2_ fold change from the average gene expression in respective control mice. Each column is one sample. Asterisks indicate FDR adjusted p-value from original sequencing data, with asterisks on the left of the gene name describing the silencing condition vs respective control and asterisks on the right of the gene name describing activating vs respective control (*p-adj<0.05; **p-adj<0.01). (B) Quantification of percentage of CD31+ vascular length that is also LDLR+ in DREADDs^activating^ mice and littermate controls 3 hours after CNO injection. Increasing neuronal activity increases percent vascular length that is LDLR+ (n=8-9; p=0.0033, unpaired two-tailed t-test). Error bars represent SEM. (C) Heat map of cholesterol gene expression in a brain endothelial stem cell model after exposure to low (L) or high (H) shear stress. Human iPSCs were differentiated into endothelial-like cells and treated with CHIR to further induce brain endothelial properties. Cells were exposed to low or high shear stress for 72 hours and RNA was isolated for sequencing. Asterisks indicate adjusted p-value (*p-adj<0.05; **p-adj<0.01; ***p-adj<0.001; ****p-adj<0.0001). Heat map color scale indicates log_2_ fold change from the average gene expression in low shear stress condition.

### Shear stress regulates brain endothelial cholesterol synthesis in vitro

To further understand how neuronal activity controls endothelial cholesterol synthesis and uptake, we wanted to know whether modulation of brain endothelial cholesterol metabolism might be downstream of increased blood flow. We differentiated human induced pluripotent stem cells (iPSCs) into endothelial-like cells and used CHIR99021 to activate Wnt signaling and further induce CNS endothelial properties. We then exposed the cells to either minimal (∼0 dyne/cm^2^) or high (∼12 dyne/cm^2^) shear stress for 72 hours before isolating RNA and performing RNA sequencing. We found that high shear stress significantly upregulates the expression of genes involved in cholesterol synthesis (*Hmgcr*, *Mvd*, *Fdps*, *Fdft1*, *Sqle*, *Cyp51*, *Msmo1*, *Nsdhl*, *Dchr7*, *Sc5d*, *Dhcr24*), synthesis regulation (*Srebf2*), and uptake (*Ldlr*) (**Fig 1C**).

As in our *in vivo* activity manipulation, expression of the cholesterol efflux transporter *Abca1* trended in the opposite direction (lower with high shear stress). These results suggest that increased blood flow might mediate the activity-dependent modulation of endothelial cholesterol synthesis and uptake.

### Brain endothelial cholesterol synthesis and uptake is not regulated by dietary fat intake

To understand what other signals might regulate cholesterol synthesis and uptake in brain endothelial cells, we investigated whether increasing dietary cholesterol levels would alter the same gene cassette. Adult wildtype mice on vivarium chow were switched to either control diet (12% kcal fat, 40 mg/kg cholesterol) or high fat diet (60% kcal fat, 280 mg/kg cholesterol) for one month before brain endothelial cells were isolated for RNA sequencing. Interestingly, high fat diet did not affect brain endothelial expression of cholesterol-related genes (**Fig 2A**). These data demonstrate that brain endothelial cholesterol metabolism is not regulated by dietary cholesterol.

**Figure 2.**
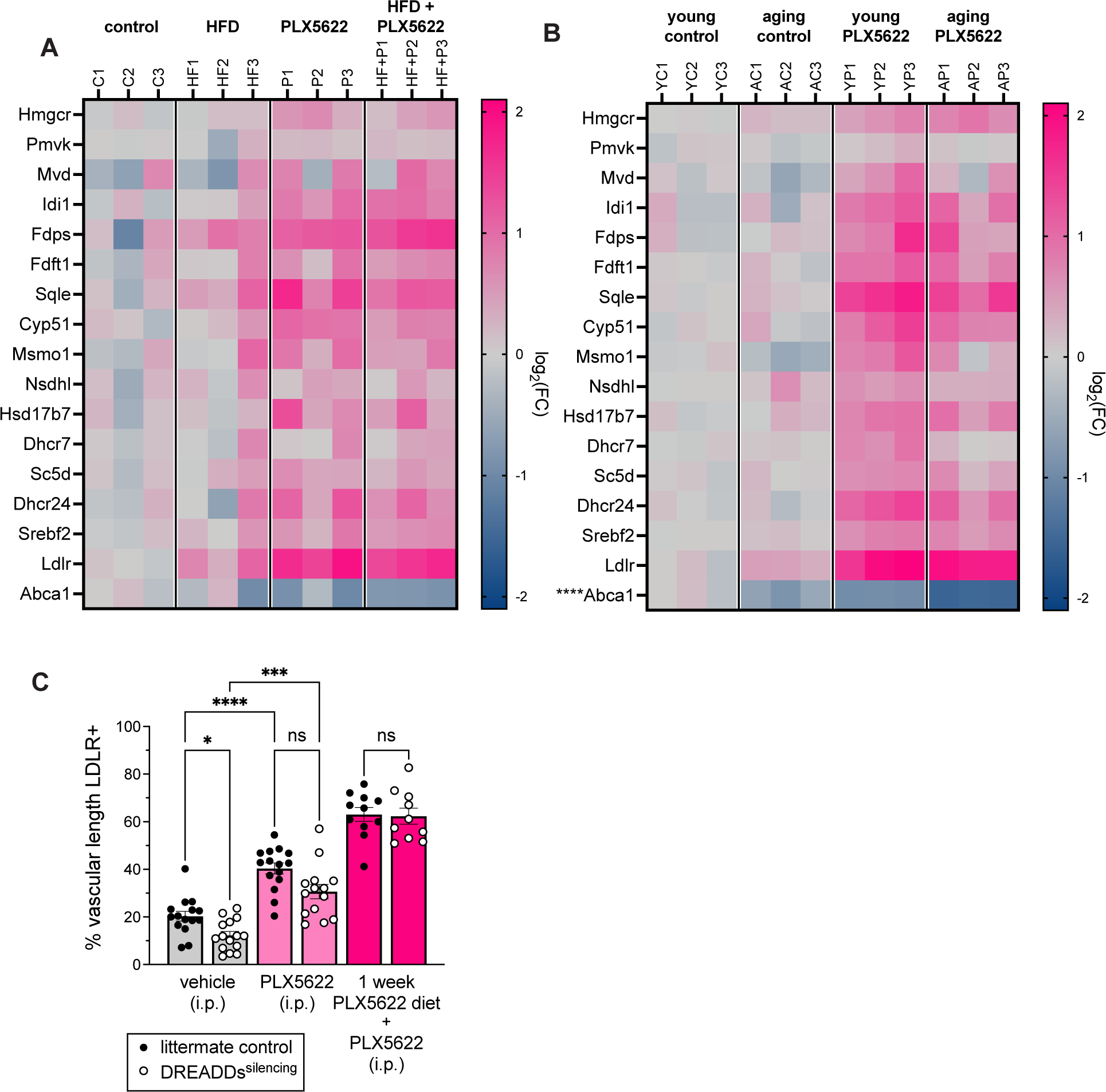
Brain endothelial cholesterol synthesis and uptake are not regulated by diet or age, and PLX5622 acts downstream of neuronal activity. (A) Heat map of cholesterol gene expression in brain endothelial cells after one month of control (C), high-fat (HF), PLX5622 (P), or high-fat+PLX5622 (HF+P) diet. Endothelial cells were isolated by FACS, and bulk RNA sequencing was performed on isolated mRNA. Each column is one sample. No cholesterol-related genes had a p-adj<0.05 in the comparison between control vs high-fat diet groups. Heat map color scale indicates log_2_ fold change from the average gene expression in mice on control diet. (B) Heat map of cholesterol gene expression in brain endothelial cells at age 12 weeks (young, Y) or 17.5 months (aging, A). Mice were raised on vivarium chow and switched to control (C) or PLX5622 (P) diet starting at 8 weeks of age to the time of tissue collection. Endothelial cells were isolated by FACS, and bulk RNA sequencing was performed on isolated mRNA. Each column is one sample. Asterisks indicate adjusted p-value (*p-adj<0.05; **p-adj<0.01; ***p-adj<0.001; ****p-adj<0.0001) for young vs aging mice on control diet. Heat map color scale indicates log_2_ fold change from the average gene expression in young mice on control diet. (C) Quantification of percentage of CD31+ vascular length that is also LDLR+ in DREADDs^silencing^ and control mice injected with PLX5622 or vehicle at the onset of the dark (waking) cycle (time=0). All mice were injected with CNO at times 0, 4, and 8 hours to maintain neuronal silencing, and tissue was collected at t=12 hours. Silencing reduced percent vascular length that was LDLR+ while PLX5622 increased it (p=0.0229; p=0.0002). There was a trend towards lower percent vascular length LDLR+ in the PLX5622+silencing group compared to the PLX5622 group (p=0.0827). Experiment was repeated in a second cohort of DREADDs^silencing^ and control mice fed PLX5622 diet for one week. After one week of PLX5622 diet, there was no difference between control and silencing groups (p>0.999). n=10-15 per group; Brown-Forsythe and Welch ANOVA. Error bars represent SEM.

### Brain endothelial cell cholesterol metabolism does not change with age

Astrocytes, one of the major producers of cholesterol in the brain, show decreased expression of cholesterol synthesis enzymes in the aged brain^25^. We therefore wanted to understand whether brain endothelial cell cholesterol metabolism also changes with age. We performed RNA sequencing on brain endothelial cells isolated from 3-month-old and 17.5-month-old mice. Expression of cholesterol synthesis enzymes and uptake receptor did not significantly change with age. Expression of the cholesterol efflux transporter (*Abca1*) was slightly decreased with age (−0.37 fold change; p-adj <0.0001) (**Fig 2B**). These data show that, overall, expression of cholesterol synthesis machinery in brain endothelial cells is remarkably stable with age, unlike in astrocytes.

### PLX5622 acts downstream of activity in regulating brain endothelial cholesterol synthesis and uptake

Previously, we found that PLX5622 treatment increases brain endothelial expression of genes involved in cholesterol synthesis and uptake, and this effect was independent of microglial depletion^23^. We thus also included PLX5622 diet in our experiments assessing the contribution of diet and age to regulation of brain endothelial cholesterol pathways. Neither dietary fat nor age interacted with PLX5622 in regulating the expression of cholesterol-related genes in brain endothelial cells (**Fig 2A-B**).

To understand how PLX5622 treatment and neuronal activity interact in the context of brain vascular cholesterol synthesis and uptake, we performed an epistasis experiment in which we combined PLX5622 treatment (which increases brain endothelial cholesterol gene expression) and silencing neuronal activity (which decreases brain endothelial cholesterol gene expression). Specifically, *CamKII*α*-*tTA; TRE-hM4Di (DREADDs^silencing^) mice and littermate controls were injected with PLX5622 (50 mg/kg) or vehicle at the beginning of the dark cycle (awake period). They were also injected with CNO at the onset of the experiment and every four hours throughout the 12-hour dark cycle. At the end of the dark period, mice were perfused, and their brains were analyzed for vascular LDLR. As expected, silencing neuronal activity decreased vascular LDLR, and PLX5622 injection increased it (p=0.0229; p=0.0002) (**Fig 2C**). While not significant, there was a trend towards lower vascular LDLR in the PLX5622+silencing group compared to the PLX5622 group, with silencing activity decreasing vascular LDLR by ∼10% (p=0.0827) (**Fig 2C**). However, when we performed this experiment with mice that were fed PLX5622 diet for one week prior to CNO injections, there was no longer even a trend towards LDLR decrease after neuronal silencing (p>0.999) (**Fig 2C**). These data suggest that PLX5622 is acting downstream of neuronal activity in regulating brain endothelial cholesterol metabolism. If PLX5622 were acting upstream of neuronal activity, we would still expect to see a decrease in LDLR after negatively modulating neuronal activity in PLX5622-treated mice. Given our findings that PLX5622 affects cholesterol pathways only in CNS endothelial cells, and not in other brain cell types or in peripheral endothelial cells^23^, we used PLX5622 to pharmacologically target CNS endothelial cholesterol synthesis and uptake pathways to probe the physiological consequences of modulating brain endothelial cholesterol metabolism.

### Increasing brain endothelial cholesterol synthesis and uptake does not alter brain cholesterol levels

The current dogma is that peripheral and brain cholesterol are largely separate pools, and the brain synthesizes its own cholesterol, with most synthesis occurring in astrocytes and oligodendrocytes^26–28^. Thus, we aimed to understand whether an increase in brain endothelial cell cholesterol metabolism might serve to boost overall brain cholesterol content. We thought it unlikely that endothelial cells would contribute to brain cholesterol levels, because even the highest observed expression levels of synthesis enzymes in endothelial cells are dwarfed by expression levels of the same genes in astrocytes and oligodendrocytes^29^. To test whether increasing brain endothelial cholesterol metabolism affects brain or peripheral cholesterol levels, we isolated forebrain, liver, and serum samples from adult wildtype mice fed control or PLX5622 diet (which upregulates brain endothelial cholesterol synthesis and uptake) for one month and performed lipidomics analysis via mass spectrometry. There were no significant differences in cholesterol concentration in forebrain, liver, or serum samples from male or female mice (**Fig 3A**). These data suggest that changes in brain endothelial cholesterol metabolism serve a more local function within endothelial cells.

**Figure 3.**
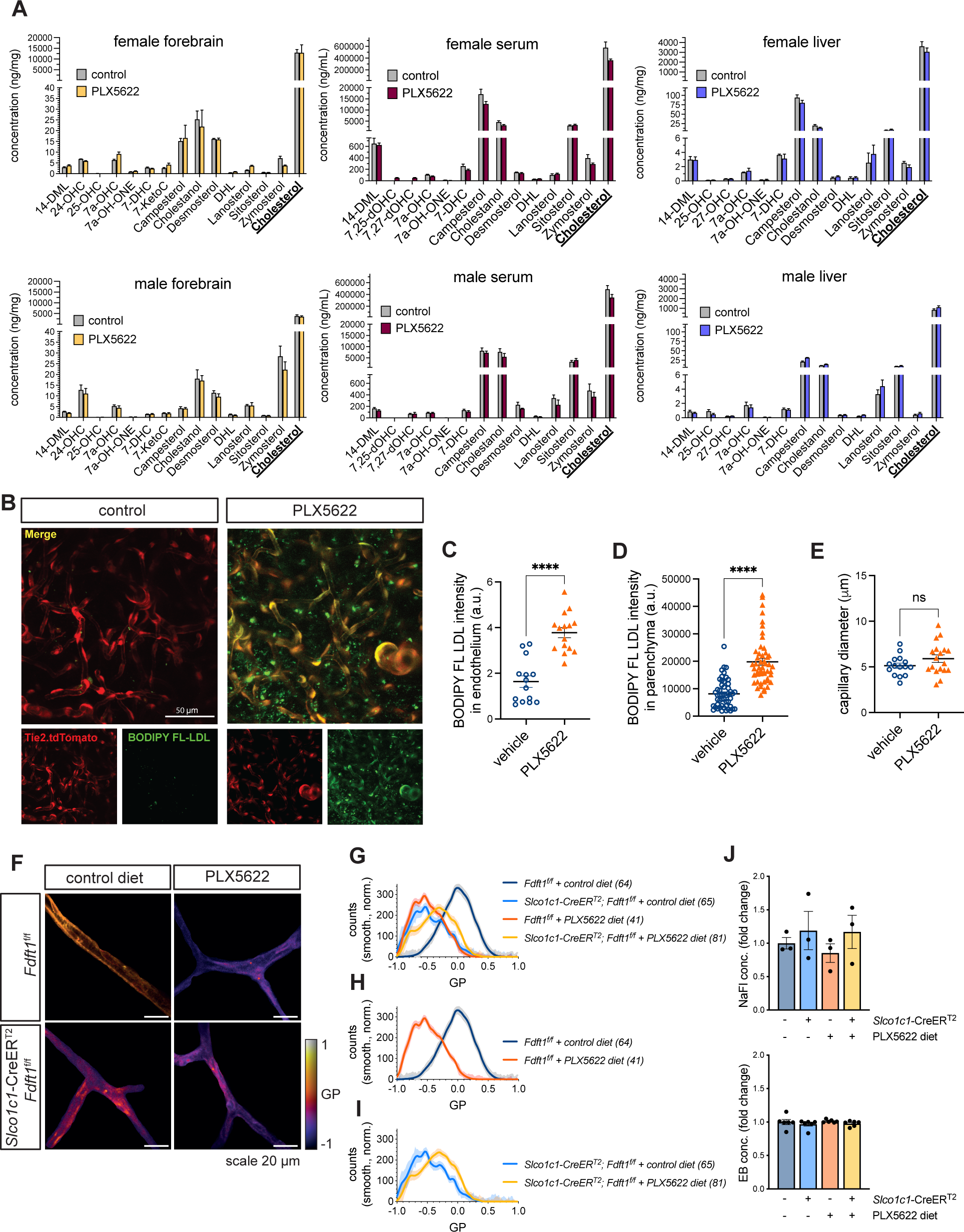
Increasing expression of brain endothelial cholesterol synthesis and uptake genes does not alter brain cholesterol content but increases vascular LDL uptake and alters endothelial membrane lipid order. (A) Lipidomics analysis of serum, forebrain, and liver from adult male and female mice fed control or PLX5622 diet for one month. There were no significant differences between diet groups for any detected sterol. n=3 per group; Mann-Whitney test. Error bars represent SEM. (B) Representative images from 2-photon visualization of LDL uptake approximately 90 min after BODIPY FL LDL injection. *Tie2*-Cre; *Ai14*^f/f^ mice were injected i.p. with PLX5622 (50mg/kg) or vehicle 24 hours before imaging. Five minutes prior to imaging, mice were injected retro-orbitally with BODIPY^TM^ FL LDL (50 μL, 1mg/mL). PLX5622 treatment increases the amount of BODIPY^TM^ FL LDL (green) uptake in endothelial cells (red). (C) Quantification of BODIPY^TM^ FL LDL fluorescence intensity in endothelial cells 60 minutes after BODIPY^TM^ FL LDL injection. A binary mask was created after applying an intensity threshold in the red channel (endothelium). The mask was applied to the BODIPY™ FL LDL channel to measure intensity within the endothelium. Each data point represents one image stack. Stacks were obtained across three mice per condition. PLX5622 increases LDL uptake in endothelial cells. n=14-15 stacks; p<0.0001, unpaired t-test. Error bars represent SEM. (D) Quantification of BODIPY^TM^ FL LDL brain parenchyma fluorescence intensity within carefully placed, hand-drawn ROIs in image stacks from 60-90 minutes after BODIPY^TM^ FL LDL injection. Each data point represents one ROI. ROIs were obtained across three mice per condition. PLX5622 increases BODIPY signal in the parenchyma. n=50 ROIs; p<0.0001, Mann-Whitney test. Error bars represent SEM. (E) Quantification of capillary diameter in *Tie2*-Cre; *Ai14*^f/f^ mice injected with PLX5622 (50mg/kg) or vehicle 24 hours before imaging. There was no significant difference in baseline capillary diameter. n=15-18, p=0.1457, unpaired t-test. Error bars represent SEM. (F) Representative images of vasculature after perfusion with Laurdan dye. Color scale represents general polarization. *Slco1c1*-CreER^T^^2^; *Fdft1*^f/f^ and *Fdft1*^f/f^ littermate controls were fed control or PLX5622 diet for one month. Laurdan dye was perfused through vasculature and its intercalation into the vascular membrane was assessed with 2-photon imaging. (G) General polarization shift of all groups described in (F). (H) General polarization shift in *Fdft1*^f/f^ mice on control and PLX5622 diet (I) General polarization shift in *Slco1c1*-CreER^T^^2^; *Fdft1*^f/f^ mice on control and PLX5622 diet (J) Quantification of BBB permeability to sodium fluorescein and Evans blue in *Slco1c1*-CreER^T^^2^; *Fdft1*^f/f^ and *Fdft1*^f/f^ littermates after one month of control or PLX5622 diet. Tracers were injected i.v. and allowed to circulate 4 hours before perfusion. Tracers were extracted and concentration was calculated using a standard curve from tracer-spiked brain samples. BBB permeability was not altered by FDFT1 ECKO, PLX5622 diet, or the combination. n=3 per group. Error bars represent SEM.

### PLX5622 increases LDL uptake in brain endothelial cells

We have found that both increasing neuronal activity and PLX5622 treatment increases brain endothelial expression of LDLR (**Fig 1A**)^23^. Here we aimed to use PLX5622 to understand whether this increase in CNS endothelial cell expression of LDLR translates to a functional increase in endothelial LDL uptake from the blood. We injected mice intraperitonially with PLX5622 (50 mg/kg) or vehicle, and 24 hours later injected the same mice retro-orbitally with a fluorescently labeled BODIPY^TM^ FL LDL molecule. We performed *in vivo* 2-photon imaging of brain vasculature and observed a stark increase in the amount of BODIPY^TM^ FL LDL present in the vasculature after PLX5622 injection (p<0.0001) (**Fig 3B-C**). We also found a significant increase in parenchymal BODIPY signal (p<0.0001) (**Fig 3B,D).** There was no change in resting capillary diameter between the two groups (**Fig 3E**). Together, these data clearly demonstrate that increased expression of endothelial LDLR causes a functional increase in brain vascular LDL uptake.

### Increasing brain endothelial cholesterol synthesis and uptake alters endothelial membrane properties

We have shown that, in response to neuronal activity, shear stress, and PLX5622 treatment, endothelial cells upregulate cholesterol synthesis and uptake machinery while downregulating cholesterol efflux, and we have shown that this leads to increased endothelial LDL uptake. We hypothesized that these changes in cholesterol metabolism and uptake might alter endothelial membrane organization. To understand whether PLX5622 treatment alters the biophysical properties of the endothelial membrane, we stained the vasculature with Laurdan dye, which intercalates into cell membranes differently based on the composition of the phospholipid bilayer. We found that one month of PLX5622 treatment caused a clear shift in general polarization (GP) of the dye in brain vasculature (**Fig 3F, H**). To confirm that this shift was connected to changes in cholesterol metabolism, we also tested mice with brain endothelial knockout (ECKO) of farnesyl-diphosphate farnesyltransferase (*Fdft1*), an enzyme in the cholesterol synthesis pathway. As expected, on control diet, *Fdft1* ECKO mice also displayed a GP shift compared to littermate controls (**Fig 3F-G**). Interestingly, the effect of PLX5622 on GP was greatly reduced in *Fdft1* ECKO mice compared to littermate controls (**Fig 3 F-G, I**). These data suggest that PLX5622 indeed causes biophysical changes in the endothelial membrane and that these changes are in part dependent on the cholesterol synthesis pathway.

We also tested BBB permeability in these mice to confirm our previous findings that PLX5622 does not alter endothelial barrier permeability to tracers^23^. We found that neither PLX5622 diet nor *Fdft1* ECKO nor the combination altered BBB permeability to sodium fluorescein or Evans blue (**Fig 3J**).

### Endothelial cholesterol inhibits arteriole dilation in response to capillary K^+^ stimulation

We have shown so far that neuronal activity and shear stress regulate brain endothelial cholesterol synthesis and uptake. We have also demonstrated that, while changes in endothelial cholesterol pathways do not alter overall brain cholesterol levels, they can modify endothelial membrane biophysics. Given these results and previous reports that cholesterol can modulate endothelial K^+^ signaling^17–21^, we hypothesized that endothelial cholesterol might be dynamically regulated as a part of NVC. Neuronal activity-generated extracellular K^+^ can activate capillary K_IR_2.1, causing a hyperpolarizing current that travels upstream, leading to arteriole dilation and increased blood flow^13^. We tested whether PLX5622-induced increases in cholesterol synthesis and uptake would affect this retrograde signaling. Specifically, mice were fed control or PLX5622 diet for at least one month, and the *ex vivo* capillary-parenchymal arteriole (CaPA) preparation^13,30,31^ was used to measure arteriole dilation in response to capillary K^+^-stimulation. This *ex vivo* preparation (which contains just vascular cell types) was particularly ideal in that it removed any confounding factors of microglial depletion or changes in baseline neuronal activity due to PLX5622, focusing only on vascular physiology. A glass pipette was used to stimulate either the capillaries or arteriole of isolated vascular segments (**Fig 4A**). Remarkably, PLX5622 diet abolished arteriole dilation in response to capillary K^+^ stimulation (**Fig 4B-C**). Direct arteriole stimulation with K^+^ was still able to generate a dilation response (**Fig 4B**), showing that the arteriole itself was still capable of dilation. As PLX5622 causes increased expression of cholesterol synthesis and uptake genes throughout the vascular tree^23^, this preserved response is likely occurring through K_IR_ channels in the SMCs wrapped around the arterioles. These results demonstrate that PLX5622 causes a physiological change in brain vasculature independent of changes in baseline neuronal activity or presence of microglia. This K^+^ response inhibition might contribute to NVC deficits previously reported in mice fed PLX3397, a similar, less specific CSF1R inhibitor^32^.

**Figure 4.**
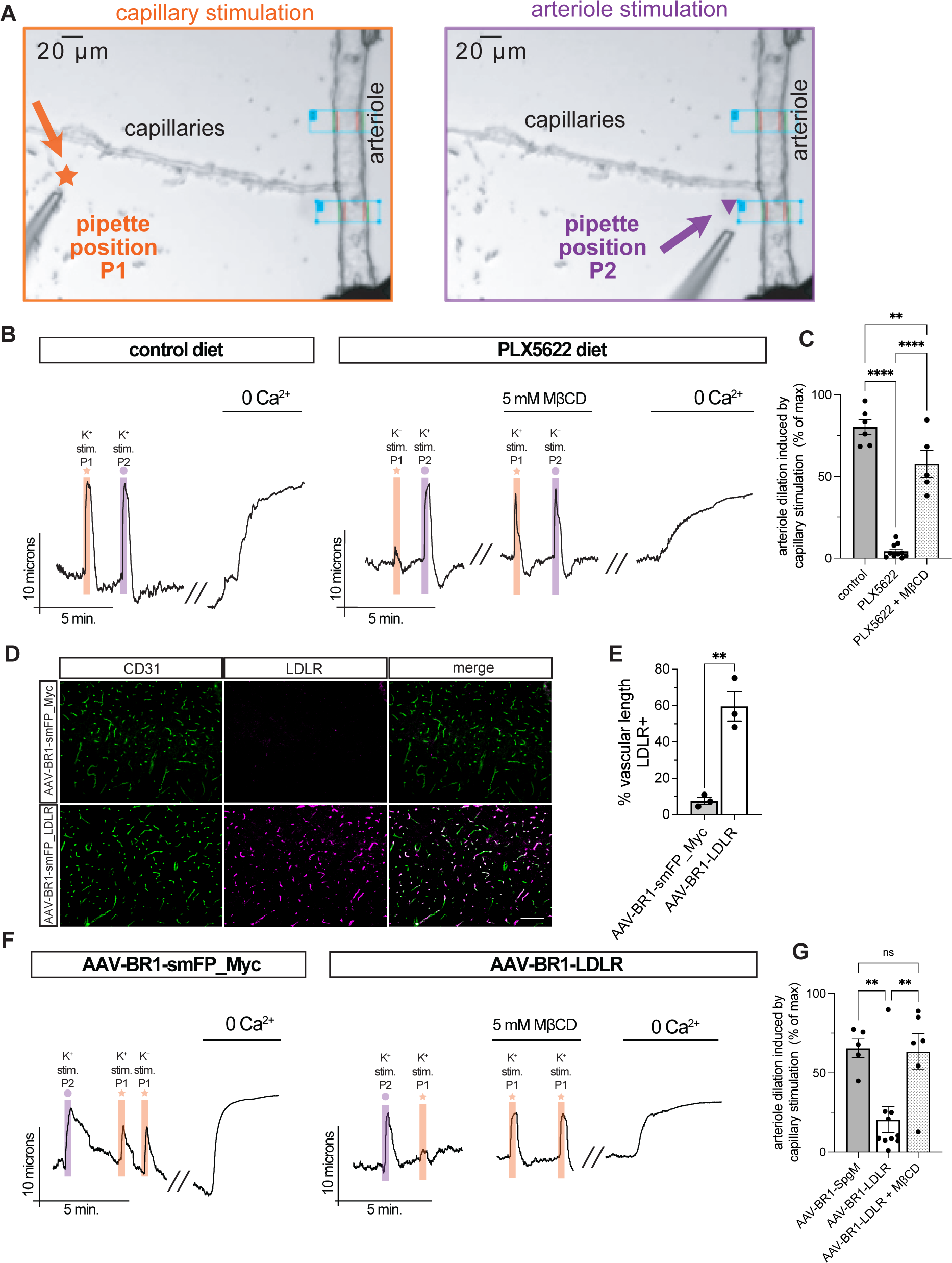
Endothelial cholesterol inhibits arteriole dilation in response to capillary K^+^ stimulation. Mice were euthanized with sodium pentobarbital and decapitated. Vascular segments containing both capillaries and arterioles were dissected from cortical tissue surrounding the middle cerebral artery. The arteriole was cannulated on one end and tied off on the other to maintain pressurization. (A) Representative images of pipette placement for K^+^ stimulation of capillaries (left, orange) and arterioles (right, purple) (B) Representative traces of arteriole dilation after K^+^ stimulation of capillaries and arterioles from mice fed control (left) or PLX5622 diet (right) for at least 1 month before ex vivo preparation. PLX5622-mediated deficits in arteriole dilation following capillary stimulation were rescued by cholesterol depletion with 30-minute bath application of 5mM MβCD. Bath application of 0 Ca^2+^ demonstrates maximum dilation capacity of arterioles. Scale bars represent 10 microns dilation and 5 minutes time. (C) Quantification of experiment shown in (B). Arteriole dilation in response to K^+^ stimulation is represented as percentage of maximum dilation. PLX5622 induced deficits in dilation response (p-adj<0.0001) were rescued by MβCD (p-adj<0.0001). Rescued response was still significantly lower than control response (p-adj=0.0099). n=5-10, one-way ANOVA with Bonferroni’s multiple comparisons test. Error bars represent SEM. (D) Representative images of cortical sections from mice injected with AAV-BR1-smFP_Myc or AAV-BR1-LDLR. Sections stained with antibodies against CD31 (green) and LDLR (magenta). Scale bar represents 100 μm. (E) Quantification of percentage CD31+ vascular length that is also LDLR+. AAV-BR1-LDLR significantly increases vascular LDLR. n=3; p=0.0033, unpaired t-test. (F) Representative traces of arteriole dilation after K^+^ stimulation of capillaries and arterioles from mice injected with AAV-BR1-smFP_Myc (left) or AAV-BR1-LDLR (right) 2-4 weeks before ex vivo preparation. Brain endothelial cell LDLR overexpression-mediated deficits in arteriole dilation following capillary stimulation were rescued by cholesterol depletion with 30-minute bath application of 5mM MβCD. Bath application of 0 Ca^2+^ demonstrates maximum dilation capacity of arterioles. Scale bars represent 10 microns dilation and 5 minutes time. (G) Quantification of experiment shown in (D). Arteriole dilation in response to K^+^ stimulation is represented as percentage of maximum dilation. LDLR overexpression in brain endothelial cells induced deficits in dilation response (p-adj=0.0097) were rescued by MβCD (p-adj=0.0088). Rescued response was not significantly different than control response (p-adj>0.9999). n=5-10, one-way ANOVA with Bonferroni’s multiple comparisons test. Error bars represent SEM.

We hypothesized that PLX5622 abolishes retrograde hyperpolarization because higher levels of endothelial cholesterol act to inhibit endothelial K_IR_ channels. We therefore tested whether depleting cholesterol from PLX5622 vessels would rescue arteriole dilation response. We used methyl-beta-cyclodextrin (MβCD) to deplete membrane cholesterol from the vessels of PLX5622-treated mice. Cholesterol depletion significantly rescued arteriole dilation in response to capillary K^+^ stimulation (**Fig 4B-C).** These results suggest that increased endothelial cholesterol contribute to PLX5622-induced deficits in endothelial retrograde signaling. Together, these data lead us to hypothesize that neuronal activity increases endothelial cell cholesterol synthesis and uptake, and this cholesterol acts to inhibit activity-induced vasodilation, acting as a negative feedback mechanism for hyperemia.

PLX5622 increases brain endothelial expression of both cholesterol synthesis enzymes and the cholesterol uptake receptor. To test whether increase of just cholesterol uptake in brain endothelial cells would be sufficient to inhibit arteriole dilation, we created an AAV-BR1 virus to overexpress LDLR in brain endothelial cells^33^. As a control, we made an AAV-BR1 virus expressing spaghetti monster fluorescent protein with a myc tag (smFP_Myc). The AAV-BR1-LDLR virus, but not the AAV-BR1-smFP_Myc virus, caused a stark increase in CD31+ vascular length also positive for LDLR (**Fig 4D-E)**. We then repeated the CaPA preparation and K^+^ stimulation in mice injected with AAV-BR1-LDLR or AAV-BR1-smFP_Myc. Compared to mice injected with the control virus, mice overexpressing LDLR in brain endothelial cells exhibited inhibited arteriole dilation in response to capillary K^+^ stimulation. Arteriole dilation in response to arteriole K^+^ stimulation was preserved (**Fig 4F),** likely indicating that SMCs were unaffected by the virus and still able to relax in response to direct K^+^ stimulation. To confirm that LDLR-mediated inhibition of arteriole response was related to cholesterol, we tested whether bath application of MβCD was sufficient to rescue the deficit. We found that indeed MβCD rescued arteriole dilation in response to capillary K^+^ stimulation (**Fig 4F-G**). Overall, these results suggest that increased endothelial cholesterol from either PLX5622 treatment or LDLR overexpression inhibits a key aspect of neurovascular coupling: the ability of capillaries to sense K^+^ and send a retrograde signal to upstream vasculature to elicit vasodilation. We thus hypothesize that neuronal activity dynamically regulates CNS endothelial cholesterol metabolism as a negative feedback mechanism for NVC.

## Discussion

Here we find that CNS endothelial cholesterol synthesis and uptake is regulated by neuronal activity, with neuronal activation leading to increases in endothelial cholesterol synthesis and uptake. We show that upregulation of brain endothelial cholesterol synthesis, either by PLX5622 administration or virus-mediated LDLR overexpression, inhibits retrograde hyperpolarization and arteriole dilation in response to capillary K^+^ stimulation. In both cases, this deficit in K^+^-mediated arteriole dilation is rescued by depletion of membrane cholesterol. Together, these data suggest that increases in endothelial cholesterol act as a negative feedback mechanism in neurovascular coupling.

While several studies provide evidence that K_IR_ channels are cholesterol-sensitive^20,21,34–36^, and while we show that endothelial response to K^+^ is inhibited by cholesterol, we cannot say whether cholesterol inhibits endothelial retrograde signaling solely by affecting K_IR_ channels. Cholesterol might have a myriad of effects on cell signaling processes in the cell. In the future it will be important to understand the range of functional effects of increased endothelial cholesterol content.

Functional hyperemia occurs in a matter of seconds after neuronal activity and is transient in nature. What we have captured in our transcriptional data is an endothelial response to sustained neuronal activity lasting over an hour^24^. We also see an increase in cholesterol-related gene expression three hours after kainic acid-induced seizures^37^. From these data, it is evident that sustained neuronal activity leads to transcriptional changes in a cholesterol-related gene cassette. However, it remains unclear whether cellular cholesterol stores or cholesterol uptake are utilized on much shorter timescale to help vasodilation return to baseline following hyperemia. Alternatively, it may be that endothelial cholesterol metabolism provides negative feedback to NVC processes specifically in cases of sustained or pathological neuronal hyperactivity.

Interestingly, although we found our cholesterol phenotype to be independent of microglial depletion^23^, some recent papers have suggested a link between microglial depletion and neurovascular coupling. Bisht et al. show that PLX3397 treatment increases baseline cerebral blood flow (CBF) and capillary diameter, and that PLX3397-treated mice have a lower percent increase in CBF in response to CO_2_^32^. These effects were phenocopied in *P2ry12*^-/-^ and *Panx*^-/-^ mice, and the authors thus suggest that microglial-vascular purinergic signaling is involved in neurovascular coupling^32^. Császár et al. similarly find that PLX5622 treatment, pharmacological P2RY12 inhibition, and P2RY12 knockout increase baseline CBF and decrease CBF response to a variety of manipulations, including whisker stimulation and hypercapnia^38^. Given our findings, it is likely that PLX5622 or PLX3397 treatment used in these studies caused NVC deficits both via inhibition of microglia-mediated purinergic signaling as well as by altered endothelial cholesterol content. It is also possible that these studies’ manipulation of purinergic signaling with P2RY12 global knockout mice or pharmacological P2RY12 inhibition affects NVC in a third manner: P2RY12 has been found to be expressed in SMCs and play a direct role in vasodilation and constriction^39^. Overall, we do not find our results to be inconsistent with the findings of either of these papers, and in combination they help to elucidate the complex and numerous mechanisms underlying NVC.

Although we have focused on the regulation of endothelial cholesterol synthesis and uptake in healthy adult mice, the regulation of endothelial cholesterol and its effects on vascular function are also pertinent to several neuropathologies and cardiovascular diseases. Atherosclerosis is characterized by the buildup of fats—including cholesterol—on the inner walls of blood vessels, causing obstruction of blood flow. As the general dogma is that blood and brain cholesterol are separate pools, there has not been much focus on how this cholesterol might be taken up into brain endothelial cells and alter abluminal endothelial signaling. In light of the current study, it is possible that endothelial LDLR facilitates uptake of plaque-associated cholesterol, and that this uptake contributes to NVC dysfunction and cognitive deficits.

Membrane cholesterol also affects processing of amyloid precursor protein, increasing amyloid-beta (Aβ) production^40–43^. Vascular Aβ plaques, termed cerebral amyloid angiopathy (CAA), are present in up to 98% of Alzheimer’s disease (AD) patients and can also occur in the absence of AD^44^. CAA can cause stroke, dementia, inflammation, cortical microbleeds, and hemorrhage^45,46^. Despite its serious clinical ramifications, it remains unclear why plaques develop in vascular walls. One possible source is aberrant production of Aβ by endothelial cells. Interestingly, although long-term PLX5622 treatment inhibited the formation of parenchymal plaques in a mouse model of AD, Spangenberg et al. observed clear vascular plaques in PLX5622-treated mice^47^. This could be a result of dysfunctional Aβ clearance, but it could also arise from endothelial Aβ production resulting from PLX5622-mediated increases in endothelial cholesterol. Relatedly, endothelium from AD mouse models exhibits dysfunctional K_IR_ channel activity^48^, and channel function is restored by MβCD-mediated cholesterol depletion, suggesting that endothelial cholesterol contributes to K_IR_ dysfunction in AD. There is still much to uncover about changes in brain endothelial cholesterol metabolism in the context of disease.

The current study identifies neuronal activity as a modulator of CNS endothelial cholesterol synthesis and uptake and suggests a role for endothelial cholesterol in negatively regulating NVC processes. These data add to our understanding of how endothelial cells might adapt to changes in neuronal activity and play an active role in NVC. Furthermore, the results raise new questions regarding the physiological roles of brain endothelial cholesterol metabolism in health and disease.

## Acknowledgements

We would like to thank Tara Rambaldo and Neil Sekiya at the VA Flow Cytometry CORE Research Facility, Dr. Kristen Jepsen and staff at the UCSD IGM Genomics Center, and Dr. Oswald Quehenberger and staff at the UCSD LIPID MAPS Lipidomics core for their support of this work. We would like to thank Plexxikon Inc. for supplying PLX5622 for this study, Dr. Jacob Körbelin for supplying the AAV-BR1 construct, and the scientists at Virovek Inc. for optimizing our viral strategy and producing the viruses. We would also like to thank Ana Williams, John Ferrer, and UCSD Animal Care Program staff. C.P.P. was funded by an NSF GRFP, an NIH F31 (5 F31 NS110403-2), and a T32 training grant (5 T32 HL086344-13). V.C.S. was supported by a post-doctoral fellowship from the American Heart Association (20POST35160001). D.A.J. was supported by a T32 training grant (5 T32 GM 007635) and an F31 (F31HL170645). G.S. was supported by the Deutsche Forschungsgemeinschaft (SA21114/2-2) and the Wilhelm Sander-Stiftung. A.Y.S was supported by grants from the NIH/NIA (AG062738). F.D. was supported by research grants from the University of Pennsylvania Orphan Disease Center in partnership with cureCADASIL, the National Heart, Lung, and Blood Institute (R01 HL136636), and the National Institute of Neurological Disorders and Stroke (RF1 NS129022). R.D. was supported by an NIH R21 (R21 AG077148).

## Author Contributions

Conceptualization, C.P.P, R.D., F.D., and A.Y.S.; Methodology, C.P.P., K.L.F, S.P.P, and E.V.S; Validation, C.P.P., L.S., and S.A.B; Formal Analysis, C.P.P, K.L.F, L.S., V.C.S, S.A.B, and J.T.F; Investigation, C.P.P, K.L.F., L.S., V.C.S, S.A.B, J.T.F, D.A.J, and F.D.; Data Curation, C.P.P and K.L.F; Writing—Original Draft, C.P.P and R.D.; Writing—Review and Editing, C.P.P., R.D., A.Y.S, G.S., F.D., K.L.F, L.S., V.C.S., and D.A.J.; Visualization, C.P.P, L.S., V.C.S, and F.D.; Supervision, C.P.P., R.D., F.D., A.Y.S., G.S., E.V.S, and S.P.P.

## Methods

### Animals

For neuronal activity experiments, transgenic *CamKII*α*-*tTA mice^49^ were crossed with TRE-hM3Dq or TRE-hM4Di mice^50^ to produce *CamKII*α*-*tTA; TRE-hM3Dq (DREADDs^activating^) or *CamKII*α*-* tTA; TRE-hM4Di (DREADDs^silencing^) mice. For sequencing experiments, as described in the original publication^24^, DREADDs and littermate controls were injected i.p. with CNO (0.5mg/kg for DREADDs^activating^ and 1.0mg/kg for DREADDs^silencing^). Dose efficacy was confirmed with electrophysiology. Three hours after CNO injection the mice were live-decapitated using a mouse decapitator (LabScientific, XM-801) and brain endothelial cells were isolated for RNA sequencing.

For high-fat diet, aging, and lipidomics experiments, wildtype C56BL/6 mice were ordered from Charles River Laboratories or Jackson Laboratories. At 8 weeks of age, mice were switched from vivarium chow to control (D10001i), high-fat (D12492i), PLX5622 (1200 mg/kg in D10001i), or high-fat+PLX5622 (1200 mg/kg PLX5622 in D12492i) diets. All diets were purchased from Research Diets, Inc., and all were irradiated. PLX5622 was provided by Plexxikon, Inc. or purchased from MedChemExpress and incorporated into D10001 or D12492 diets by Research Diets, Inc. All experiments were approved by the Institutional Animal Care and Use Committee at UCSD.

For Laurdan, Evans blue, and sodium fluorescein experiments, *Slco1c1*-CreER^T^^2^ mice were bred to *Fdft1*^f/f^ mice^51^ at Max Planck Institute of Multidisciplinary Sciences. Eight-week-old mice were injected with 75 μg tamoxifen per g body weight for 5 days and put on control or PLX5622 (1200 mg/kg) diet 2 weeks after tamoxifen administration. Mice were kept on special diet for 4 weeks. Control and PLX5622 diets were from Research Diets, Inc., as described above. All experiments were performed in compliance with the Institutional Care and Use Committee guidelines of the Max Planck Institute and were approved by the German Federal State of Lower Saxony (Lower Saxony State Office for Consumer Protection and Food Safety).

For *in vivo* imaging experiments, *Tie2*-Cre (Jax ID: 008863) and homozygous floxed *Ai14* (Jax ID: 007914) mice from Jackson Laboratory were crossed to label the endothelium. Mice successfully expressing endothelial red fluorescence were identified using a Nightsea fluorescent protein flashlight. Experiment was performed on 4-to 10-month-old males and females, and experimental cohorts consisted of littermates matched for age and sex. A single dose of PLX5622 (Selleckchem; S8874) at 50 mg/kg, or vehicle (corn oil) was injected intraperitoneally 24 hours before imaging. All experiments were approved by the Institutional Animal Care and Use Committee at Seattle Children’s Research Institute.

For ex vivo CaPA experiments, male and female adult C57BL/6 mice ordered from The Jackson Laboratory were put on control or PLX5622 diet (Research Diets, Inc., as described above) at least one month before CaPA preparation. Viruses (1.8 x 10^11^ vg, Virovek, Inc.) were injected retro-orbitally 2-4 weeks before CaPA preparation. All experiments were approved by the Institutional Animal Care and Use Committee at University of Colorado.

### Endothelial cell enrichment

Mice were anesthetized with a cocktail of ketamine (4 mg/ml) and xylazine (0.6 mg/ml) in saline. Mice were then decapitated, brains were dissected, and cerebellum and olfactory bulbs were removed and discarded. Brains were rolled on filter paper to remove meninges. Remaining meninges and choroid plexus were removed with fine forceps. Cell preparation was performed as described in^23^. Briefly, brain tissue was diced, tissue was enzymatically digested with the Papain Dissociation System kit (Worthington, LK003176) at 35°C for 90 minutes with 95% O_2_, 5% CO_2_ continuously passed over the solution. Tissue chunks were washed with a “low-ovomucoid” EBSS solution containing 225 μg/mL ovomucoid (Worthington, LS003089). Samples were triturated with 10 mL, 5 mL, and P1000 pipette tips, successively. Cells were spun into “high-ovomucoid” solution containing 450 μg/mL ovomucoid. Cells were resuspended in a collagenase/dispase solution of 1 mg/mL collagenase type II (Worthington, LS004176) and 0.4 mg/mL neutral protease (Worthington LS02104) and incubated for 30 min at 35°C with 95% O_2_, 5% CO_2_ continuously passed over the solution. After incubation, cells were spun into high-ovomucoid solution and resuspended in a 0.5% BSA (Sigma, A4161) solution with myelin removal beads (Miltenyi Biotec, 130-096-433). After a 15 min bead incubation, 30 micron pre-separation columns (Miltenyi Biotec, 130-041-407) and LS columns (Miltenyi Biotec, 130-042-401) were used for myelin removal. Cells were blocked with rat IgG in 0.5% BSA solution on ice for 20 min. Cells were stained with AF647-conjugated anti-CD31 (Molecular Probes A14716), FITC-conjugated anti-CD13 (BD Biosciences 558744), FITC-conjugated anti-CD45 (eBioscience 11-0451-85), FITC-conjugated anti-CD11b (eBioscience), and DAPI. Cells positive for 647 and negative for FITC and DAPI were sorted into trizol at the UCSD Flow Cytometry Research Core Facility. RNA isolation was performed with the Qiagen RNeasy Micro kit (74004).

### RNA sequencing and bioinformatics (mouse)

Sequencing and bioinformatics pipeline for DREADDs mice was described in Pulido et al., 2020^24^. For high-fat diet and aging experiments, V2 non-stranded mRNA library prep and bulk sequencing were carried out by the UCSD Institute of Genomic Medicine Core. Samples were sequenced using single-read, 75 bp reads. HISAT2 was used for alignment to the GRCm38 genome, htseq-count was used to generate count tables, and differential expression was analyzed using DeSeq2. All analysis programs were run through the Galaxy open-source platform. Heat maps were generated by calculating the log_2_ of the fold change of each sample compared to the average of the control samples.

### Human iPSC differentiation and shear stress

IMR90-4 human induced pluripotent stem cells (iPSCs) were maintained in E8 stem cell maintenance media and differentiated according to the protocol described in Lian et al^52^. The CD34+ endothelial progenitor population was isolated using magnetic activated cell sorting (MACS). The resulting purified populations were frozen in liquid nitrogen in a medium containing HESFM + B27 supplemented with 30% FBS and 10% Dimethyl sulfoxide (DMSO). Cells were thawed into HESFM + B27 with FGF2 (20 ng/mL) and VEGF (20 ng/mL), as well as 4 µM CHIR99021 to induce blood-brain barrier (BBB) properties as described previously^53^. Cells were then plated into Collagen IV coated ibidi microfluidic chips at 0.8 mm and 0.6 mm heights. After 6-8 hours to facilitate attachment, chips were connected to an ibidi pump system and maintained at a flow rate sufficient to provide 0-1 dyne/cm2 to prevent starvation or oxygen deprivation for 24 hours. Pressure was increased gradually to generate either low (∼0 dyne/cm2) or high (∼12 dyne/cm 2) shear stress for 72 hours before sampling RNA or protein.

### RNA sequencing and bioinformatics (human iPSCs)

Cells were lysed and RNA collected using the Qiagen RNeasy Mini Kit. Library prep and bulk sequencing were carried out by Novagene. Samples were sequenced using paired-end reads. STAR was used for alignment to the hg38 genome, featurecounts was used to generate count tables, and differential expression was analyzed using DeSeq2 on the raw counts. Heat maps were generated from the log2 fold change and p adj vaules provided by DeSeq2.

### Immunostaining

Mice were transcardially perfused with DPBS followed by 4% paraformaldehyde in PBS. Tissue was cryopreserved in 30% sucrose, frozen in 2:1 OCT:30% sucrose and sectioned at 10 µm thickness. Sections for LDLR staining were further fixed in ice-cold methanol for 10 minutes. Sections for IBA1 staining were further fixed with 5 minutes of 4% PFA. All slides were blocked with 10% donkey serum, 0.2% triton-x-100 in PBS. Primary antibodies against LDLR (R&D Systems AF2255) and CD31 (BD Biosciences 553370) were used at 1:1000 with 10% blocking solution in PBS and incubated overnight at 4°C. Fluorescently conjugated secondary antibodies (Life Technologies) were used at 1:1000 and incubated for 1.5 hours at room temperature. Slides were coverslipped using DAPI-Fluoromount-G (SouthernBiothech 0100-20) and imaged with a Zeiss Axio Imager.D2.

### Analysis of LDLR+ vascular length

To quantify LDLR+ vascular length, sections were stained for LDLR and CD31. LDLR signal was traced using the ImageJ line tool, with each segment saved as an ROI. ROIs were then opened on the CD31 channel, and any ROI not corresponding to CD31+ vasculature was deleted. The remaining ROIs were measured, and the sum of their lengths was used as the length of LDLR+ vasculature. The rest of the LDLR-/CD31+ vascular segments were then traced in the same way, and the ROIs measured. The sum total length of all segments was used as the vascular length. Percent vascular length was calculated as (LDLR+ vessel length)/(total vessel length)*100. Statistics were performed with GraphPad Prism.

### Lipidomics

Mice were anesthetized with a cocktail of ketamine (4 mg/ml) and xylazine (0.6 mg/ml) in saline. A blood sample was collected from the right ventricle and allowed to coagulate at room temperature for serum collection. Mice were transcardially perfused with DPBS. A lobe of liver and the brain were dissected. Cerebellum, olfactory bulbs, and meninges were removed. Tissue and serum were flash-frozen. Sterol isolation, saponification, and LC-MS analysis were performed by the UCSD LIPID MAPS Lipidomics core. Samples were run on a Sciex6500 Qtrap mass spectrometer. Phenomenex Kinetics C18 2.1×150mm 1.7um columns were used. Statistics were performed with GraphPad Prism.

### Cranial window surgery

*One day prior to imaging,* anesthesia was induced with isoflurane (4% induction, 1-2% maintenance) in 100% medical oxygen. During surgery, mice were positioned on a feedback-regulated heat pad (FHC Inc.) to maintain body temperature at 37°C. To minimize disruption of the brain, thinned-skull cranial windows were generated in all animals, as described previously^54,55^.

### Two-photon microscopy

In vivo two-photon imaging system was performed with a Bruker Investigator (run by Prairie View 5.5 software) coupled to a Spectra-Physics Insight X3. Green and red fluorescence emission was collected through 525/70 nm and 615/60 nm bandpass filters, respectively, and detected by photomultiplier tubes (PMTs). High-resolution imaging was performed using a water immersion 20-X, 1.0 NA objective lens (Olympus XLUMPLFLN 20XW) using 920 nm excitation during the experiment.

### BODIPY™ FL LDL imaging and analysis

To study LDL uptake, 50 μL (1 mg/ml) of fluorescent low-density lipoprotein (BODIPY™ FL LDL; L3483; Life Technologies) was injected through the retro-orbital vein under deep isoflurane anesthesia. During imaging, isoflurane was maintained at 1.5% MAC in medical-grade air. Image stacks were collected from 10-120 minutes following injection. BODIPY™ FL LDL binding endothelium was quantified as fluorescence intensity along the capillary wall territory (endothelial Region of Interest, ROI). All image analysis was performed in in ImageJ/FIJI. For endothelial ROI selection, we created binary masks after applying an intensity threshold in the red channel. The mask was then applied to the BODIPY™ FL LDL channel to measure intensity specifically within the endothelium. LDL transport was evaluated as the extravascular/intravascular ratio of BODIPY™ FL LDL fluorescence intensity over time. Fluorescence intensity was measured within carefully placed, hand-drawn ROIs. Only raw images were utilized in data analysis, viewed with ImageJ/FIJI ver. 1.0 software. For presentation purposes, images were adjusted for contrast and cropped in Adobe Photoshop, one color channel at a time, in a similar manner across all conditions. Statistical analyses were performed using GrapPad Prism 8 software. Statistical tests and details are provided in the Figure legends. Tests of normality were first performed to validate the use of parametric tests.

### Membrane lipid order

Analysis of membrane phase properties in situ was done as described^56^. Mice were perfused to remove blood, and blood vessels were stained by perfusion with Laurdan (6-Dodecanoyl-2-Dimethylaminonaphthalene, Sigma) followed by paraformaldehyde perfusion fixation. After drop fixation overnight, brains were cut in 200 µm sagittal sections on a vibratome. Vessels were imaged by 2Photon microscopy (LaVision) and generalized polarization (GP) of vascular Laurdan fluorescence was determined as previously described^57^.

### Blood-brain barrier permeability

Measurements of BBB permeability were performed as previously described^58^. Briefly, tracers were i.v. injected (Evans blue 50 mg/g body weight; sodium fluorescein 200 mg/g body weight). After 4h incubation, mice were perfused with PBS to remove tracer from the circulatory system. The forebrain was dissected and weighed before lyophilization at a shelf temperature of –56 °C for 24h under vacuum of 0.2 mBar (Christ LMC-1 BETA 1-16) and extraction with formamide at 57°C for 24h on a shaker at 300 rpm (Eppendorf Thermomixer). Integrated density of tracer fluorescence was determined in triplicates after 1:3 ethanol dilutions. Tracer concentration was calculated using a standard curve prepared from tracer-spiked brain samples.

### AAV production

LDLR and spaghetti monster plasmids were designed by System Biosciences. The AAV-BR1 packaging vector was a gift from Jacob Korbelin. Plasmids were optimized and viruses were produced by Virovek, Inc.

### Capillary-parenchymal arteriole preparation and stimulation

Capillary-parenchymal arteriole preparation was achieved by dissecting parenchymal arterioles (and attached capillaries) arising from the middle cerebral artery as previously described^13,31^. Arteriolar segments were cannulated on glass micropipettes and the other end of the arteriole was tied off. Another glass micropipette was placed on top of the end of the capillaries to create a closed system. The vascular tree was pressurized with 40 mmHg and maintained in 37°C-heated artificial cerebrospinal fluid (aCSF): 124 mM NaCl, 3 mM KCl, 2 mM CaCl2, 2 mM MgCl2, 1.25 mM NaH2PO4, 26 mM NaHCO3 and 4 mM glucose. After the vascular segment acquired tone, 10mM K^+^ in aCSF was applied to capillaries or arterioles via pressurized ejection from a glass micropipette attached to a Picospritzer® III (Parker). IonOptix IonWizard edge detection software was used to measure arteriole lumen diameter was measured in two regions. Vessel preparations from mice on PLX5622 diet or expressing LDLR virus were treated with bath application of 5mM methyl-beta cyclodextrin (MβCD) (Cayman Chemicals 21633) for 30 minutes to deplete membrane cholesterol.

### Resource Availability

All sequencing data will be uploaded to Gene Expression Omnibus (GEO)

